# Large Organellar Changes Occur during Mild Heat Shock in Yeast

**DOI:** 10.1101/2021.01.25.428102

**Authors:** Katharina Keuenhof, Lisa Larsson Berglund, Sandra Malmgren Hill, Kara L Schneider, Per O Widlund, Thomas Nyström, Johanna L Höög

**Author notes:** Corresponding author: Johanna L Höög.

## Abstract

When the temperature is increased, the heat shock response is activated to protect the cellular environment. The transcriptomics and proteomics of this process are intensively studied, while information about how the cell responds structurally to heat stress is mostly lacking. Here, *Saccharomyces cerevisiae* were subjected to a mild continuous heat shock and intermittently cryo-immobilized for electron microscopy. Through measuring changes in all distinguishable organelle numbers, sizes, and morphologies in over 2400 electron micrographs a major restructuring of the cell’s internal architecture during the progressive heat shock was revealed. The cell grew larger but most organelles within it expanded even more. Organelles responded to heat shock at different times, both in terms of size and number, and adaptations of certain organelles’ morphology were observed. Multivesicular bodies grew to almost 170% in size, indicating a previously unknown involvement in the heat shock response. A previously undescribed electron translucent structure accumulated close to the plasma membrane during the entire time course. This all-encompassing approach provides a detailed chronological progression of organelle adaptation throughout the cellular stress response.

**Summary statement:** Exposure to mild heat shock leads to large quantifiable changes in the cellular ultrastructure of yeast, shows involvement of MVBs in the heat shock response and the apparition of novel structures.

## Introduction

Increasing the temperature activates the cell’s heat shock response. This is an ancient and evolutionarily conserved transcriptional program that results in reduced expression of genes involved in protein biosynthesis pathways and increased expression of genes encoding heat shock proteins (Verghese *et al*., 2012a). The heat shock response is also activated by other types of stressors, for example oxidative stress, exposure to heavy metals, fever, and protein conformational disorders (Morimoto and Westerheide, 2010).

Heat shock in budding yeast (*Saccharomyces cerevisiae*) is extensively used as a model to study neurodegenerative human diseases (Winderickx *et al*., 2008; Kaliszewska *et al*., 2015), where inclusion bodies of aggregated misfolded proteins accumulate at specific sites in the cytoplasm and nucleus (Takalo *et al*., 2013; Chung, Lee and Lee, 2018). Examples of such diseases are Alzheimer’s, Huntington’s, and Parkinson’s (Tenreiro *et al*., 2013).

In eukaryotes, the heat shock transcription factor (HSF) is responsible for the induction of heat shock genes. It exists in its inactive form during non-stress conditions and is activated when there is an accumulation of destabilised, misfolded proteins in the cell. HSF binds DNA and initiates transcription (Morimoto, 1993). The heat shock-induced changes in transcription and translation ensure that the cell is capable of maintaining proteostasis and metabolism when experiencing temperature stress (Mühlhofer *et al*., 2019). While many of these molecular mechanisms of the cellular heat shock response have been widely studied, the wide-reaching structural and architectural effects of such a temperature change are often neglected because attention is focused on the system of interest, e.g. the transcriptome, proteome, and/or the behaviour of misfolded proteins (Shi, Mosser and Morimoto, 1998; Verghese *et al*., 2012b).

In this study, we investigated the structural adaptations that cells undergo when subjected to a mild 38°C heat shock. A heat shock that is comparable to that used when studying protein quality control as well as temperature sensitive mutants. Using electron microscopy of high-pressure frozen cells, we obtained ultrastructural information on nearly every cellular organelle and substructure, without being limited by fluorescent labelling of a few candidate proteins such as those used in previous studies (Meaden *et al*., 1999; Lewandowska *et al*., 2006; Spokoini *et al*., 2012; Escusa-Toret, Vonk and Frydman, 2013; Meyers *et al*., 2016; Gao *et al*., 2017). To gain a nanometre-resolution map of cellular alterations prompted by heat shock that is as comprehensive as possible, a minimum of one hundred cells were imaged for each time point throughout a continuous 90 min exposure to 38°C. The time course and subsequent imaging was performed in triplicates, yielding a total of 2143 images analysed for an overview that comes as close to a screen as possible using electron microscopy. The resulting temporal map of cellular restructuring reveals that heat shock induces major morphological changes to the cells’ internal architecture and composition and that more organelles than anticipated are involved in this process.

## Results

### Heat shock has wide-reaching effects on cell structure

We used electron microscopy to study the ultrastructural changes occurring in yeast during a mild heat shock over a time course of 90 min (fig. 1A). Samples of yeast cultures were collected at 30°C and after a 5, 15, 30, 45, or 90 min shift to 38°C. These samples were cryo-immobilised using high-pressure freezing followed by freeze substitution for best possible morphological preservation (Moor, 1987; Perktold *et al*., 2007). Samples were then sectioned into 70 nm thin sections. At least 100 random images were collected at a magnification where the entire cell could be visualised in detail (9300x, pixel size 1.1 nm). The whole experiment was repeated three times.

**Figure 1.**
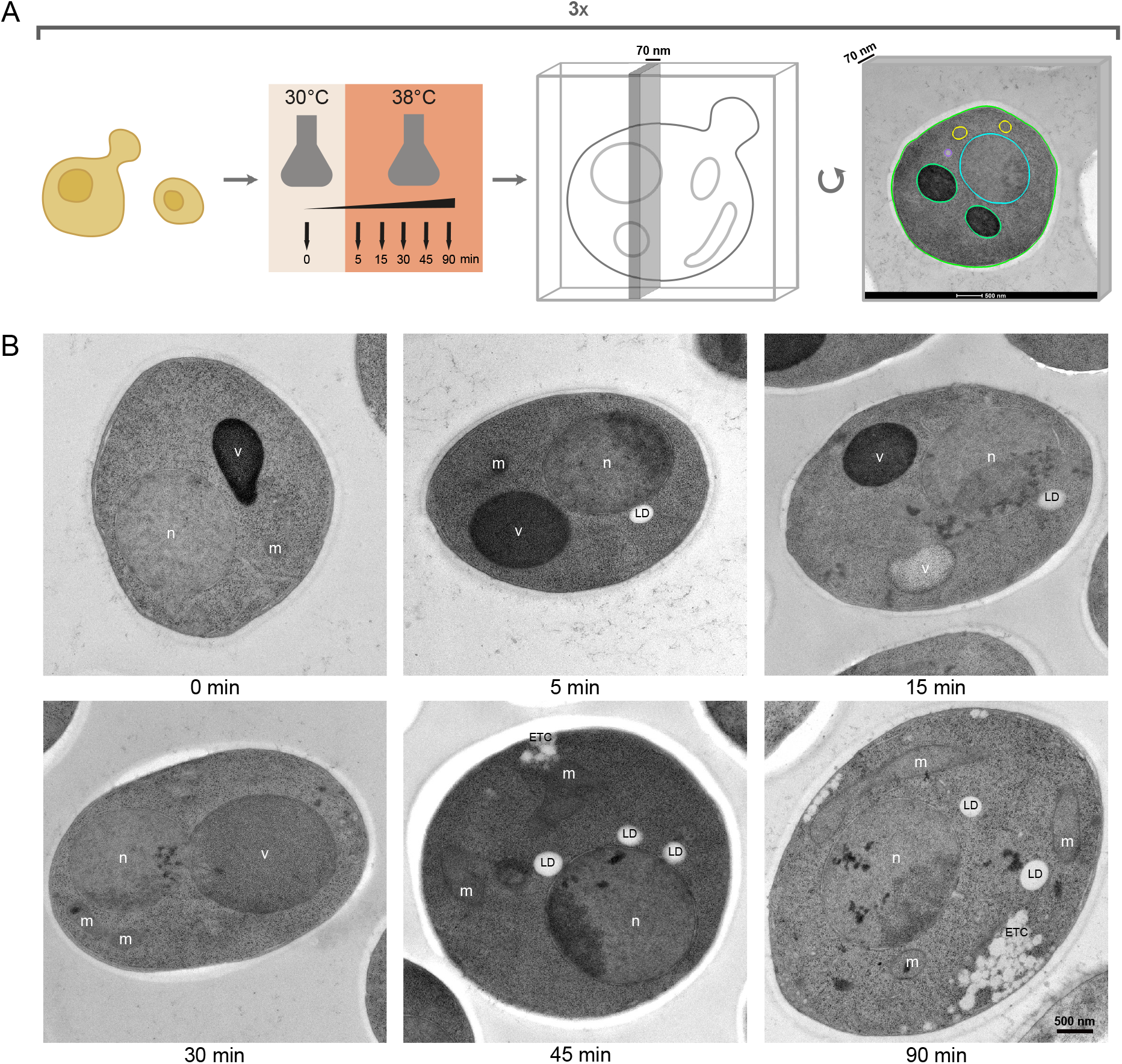
Revealing ultrastructural changes during a mild heat shock using a transmission electron microscopy screening method. A) Cartoon of the experimental procedure: Cells were grown at 30°C and heat shocked at 38°C for 0, 5, 15, 30, 45, and 90 minutes. Cells were then high-pressure frozen, freeze-substituted and sectioned to 70 nm for electron microscopy. Each heat shock experiment was done in triplicate and at least 100 images were collected of each sample, resulting in a total of 2143 images analysed. Size and morphology of all clearly visible organelles were quantified to reveal known and unknown effects of exposing yeast cells to heat shock. B) Example images show that large morphological changes and cellular reorganization occurred during heat shock. Further, a new phenotype that we call electron translucent clusters (etc) appear during heat shock. Nucleus (n), vacuole (v), mitochondrion (m), lipid droplet (ld).

Observations of cells subjected to such temperature stress revealed unexpectedly large changes in cellular architecture (fig. 1B). Organelles, such as lipid droplets (LDs) seemed to increase both in numbers and size (fig. 1B, see 5 and 45 min). Electron dense content, presumably protein aggregates, was observed in both the nucleus and mitochondria (fig. 1B, see 15, 30, 45, and 90 min). Previously undescribed electron translucent clusters (ETC) appeared often in close proximity to the plasma membrane (fig. 1B, see 45 and 90 min). However, the most prominent difference being an altered morphology of the vacuole. In heat treated cells, the vacuole appeared to change both in texture, electron density and size (fig. 1B, compare 0 and 30 min; fig. S1). To further investigate these cellular adaptations, we progressed to outline all distinguishable organelles and quantify their number and area.

### Vacuoles and the cell as a whole increase in size throughout heat shock

Outlining 2143 plasma membranes over the 6 different time points revealed that the average cell area increased by 19% over the 90 min heat shock (fig. 2A-B). A notable growth occurred between 15 and 30 min, as the first three timepoints all differ significantly from the last three.

**Figure 2.**
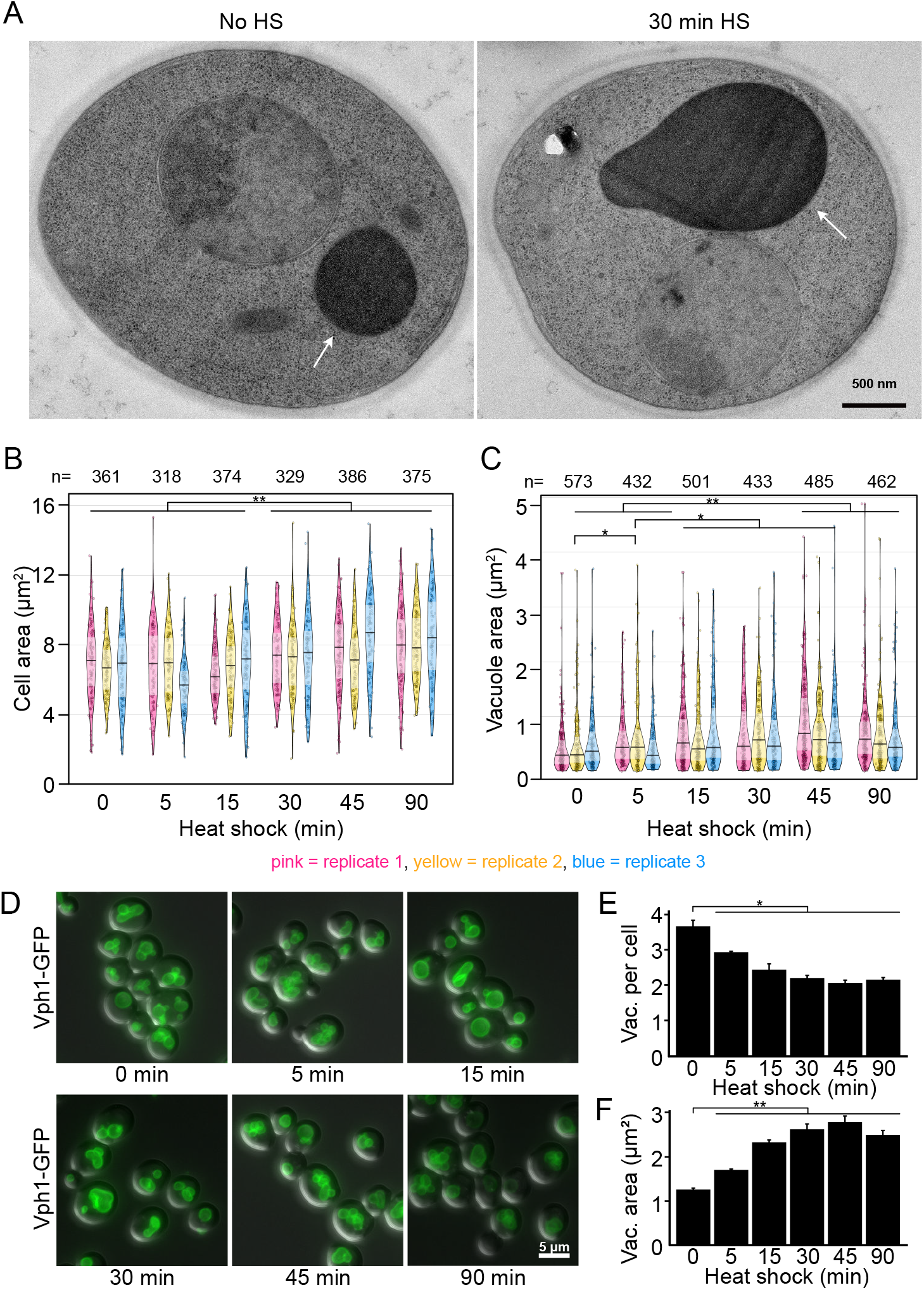
Cells and vacuoles increase in size. A) Cells with vacuoles (arrow) before and after 30 min of heat shock. B) Cell area in electron micrographs of thin sections. On average, the cell size increases by 19% over the course of 90 minutes. The pink, yellow and blue shapes represent the three separate replicates. The width represents the number of measurements within a certain range of values, one point represents one measurement. Solid line is the median and inference bands are the interquartile ranges. C) Vacuoles in electron micrographs of thin sections. On average, the vacuole size increases by 48% over the course of 90 minutes. D) Cells expressing Vph1-GFP heat shocked at indicated time points. E) Quantification of vacuole number per cell from cells in A, n > 200 per replicate and time point. F) Quantification of vacuole size from cells in A, n > 200 per replicate and time point. Bars show the mean of three replicates, error bars are the standard error of the mean. Significances: (*) p < 0.05, (**) p ≤ 0.01.

The vacuole often occupies the most amount of space in the cell, but during heat shock it grew even more compared to the cell volume (n=2866; fig. 2C). Vacuoles increased in size, peaking after 45 min of heat shock at an area 69% larger than their original size (fig. 2C). After this, they shrunk slightly but by 90 min still remained 48% larger than before heat shock.

Since electron microscopy sections are too thin to contain a whole vacuole, and may not accurately represent the number of large organelles in the cell, we also investigated their change in terms of numbers and shape throughout stress using fluorescence microscopy. Cells expressing Vph1-GFP (fig. 2D), a subunit of the vacuolar ATPase V_0_ present in the vacuolar membrane (Hecht, O’Donnell and Brodsky, 2014) were used to this purpose. Whilst the number of vacuoles decreased by 41% over the course of 90 min (non-heat shocked 1.26 ± 0.05 μm^2^, n=614, 90 min heat shock 2.49 ± 0.18 μm^2^, n=624; fig. 2E), at 45 min of heat shock they were 2.2 times as large as the control. After 90 min the vacuole was still twice as large as before heat shock (fig. 2F). This shows that vacuoles probably respond to heat shock by fusing and creating few large organelles rather than many smaller vacuoles. Further, a change to their internal morphology was also observed.

### Vacuolar texture and electron density change in response to heat shock

A striking variation in electron density of the vacuolar lumen was observed with electron microscopy. In logarithmically growing non-stressed yeast cells, vacuoles are often very electron dense. After heat shock, the vacuoles were oftentimes electron translucent and sometimes contained a grainy texture. Vacuoles were therefore classified into four major categories according to their appearance: dense, grainy, medium, and translucent (fig. 3A). The electron density of vacuoles fluctuated strongly throughout the time course in each experiment, showing lower electron densities when exposed to stress and a recovery to darkly stained vacuoles at 90 minutes (fig. 3B).

**Figure 3.**
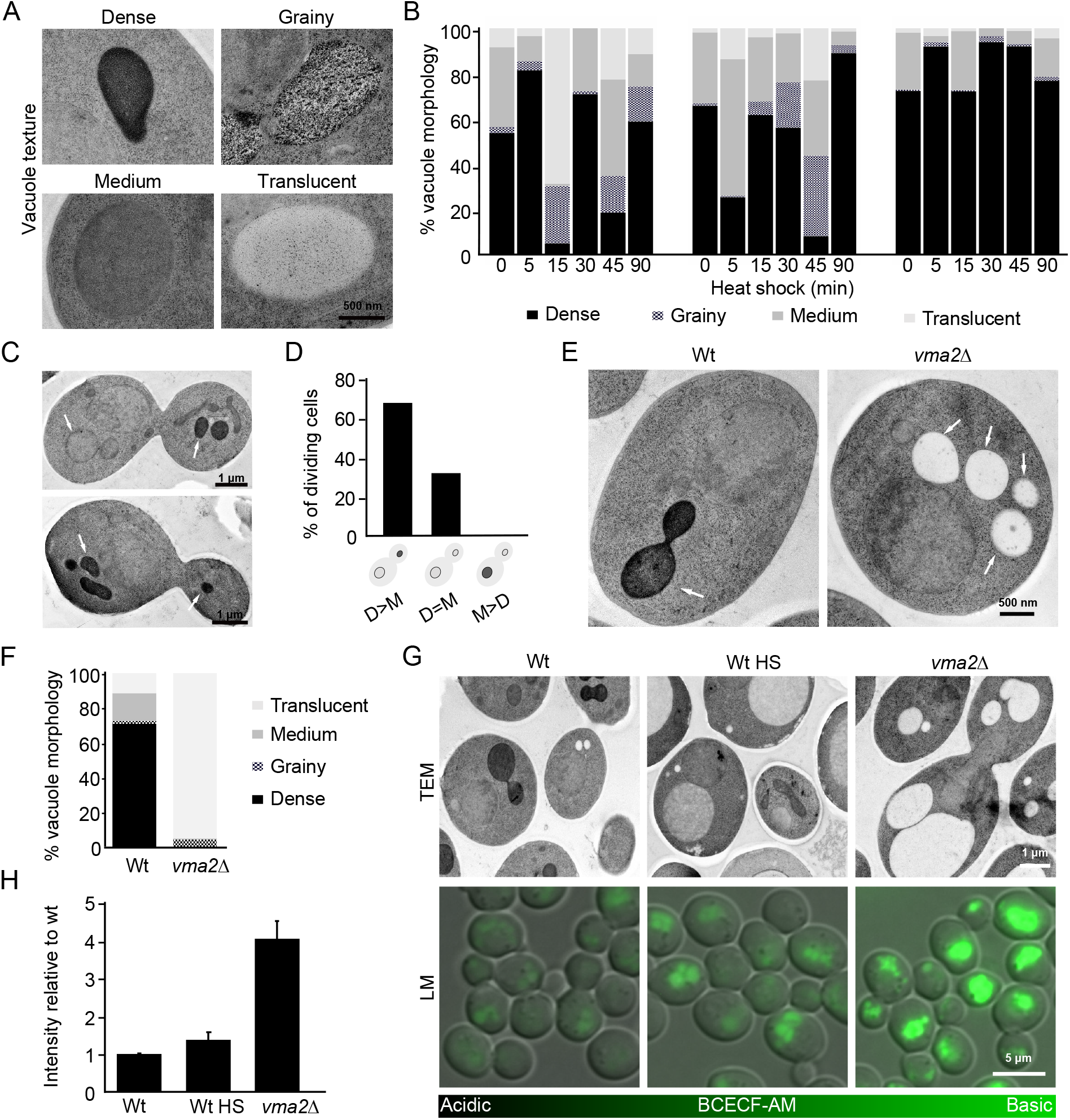
Heat shock affects vacuolar morphology and acidity. A) Gallery of different vacuolar electron densities. B) Quantification of vacuolar electron density during heat shock. More than 100 vacuoles were analyzed from three independent experiments respectively. C) EM images of dividing wt cells grown at 30°C. D) Quantification of electron density of vacuoles in mothers (M) and their daughters (D) during cell division, n = 53 dividing cells. E) EM images of Wt and *vma2*Δ grown at 30°C. F) Quantification of vacuolar electron density of E, n = 181 (Wt) and 274 (*vma2*Δ) vacuoles from a minimum of 100 cells. G) EM and fluorescence microscopy of indicated strains. HS is 45 min at 38°C. The fluorescent pH-sensitive probe BCECF accumulates in yeast vacuoles and shows vacuolar pH. The more fluorescence, the more basic vacuolar pH. H) Quantification of vacuolar BCECF fluorescence mean intensity. More than 30 vacuoles were analyzed from three independent experiments respectively. Graph shows mean relative values to wt fluorescence mean intensity with error bars corresponding to standard deviation.

When comparing electron micrographs of dividing cells, daughter cells had either a higher or equal vacuolar electron density when compared to their mother (fig. 3C and 3D). A previous study showed that upon cell division in yeast, the daughter cell has a more acidic vacuole than its older mother cell (Hughes and Gottschling, 2012). We thus hypothesized that lower electron density in vacuoles could be due to a deacidification of that organelle. The effect of vacuolar pH on the electron density of the samples was investigated using cells from a *vma2Δ* strain (fig. 3E). Vma2 is a subunit of the V1 domain of the vacuolar H^+^ ATPase and its deletion causes an increased vacuolar pH compared to its usual pH of 5-6.5 (Li and Kane, 2009). As opposed to wild type cells, almost all vacuoles in *vma2*Δ cells displayed the electron translucent phenotype, demonstrating that increased pH leads to altered vacuolar staining (fig. 3E and 3F). To determine if vacuoles deacidify during heat shock we used the pH sensitive vacuolar probe BCECF-AM, which displays increased fluorescence at increased pH (Plant *et al*., 1999; Hughes and Gottschling, 2012). As a positive deacidification control, BCECF-AM staining also confirmed the expected elevated pH in the *vma2Δ* mutant (fig. 3G and 3H). Vacuoles exposed to 45 min heat shock fluoresced brightly, compared to those of cells grown at 30°C, confirming that heat shock causes elevated internal pH in vacuoles (fig. 3G and 3H). Therefore, using our protocol, changes in vacuolar pH can be observed as altered electron density of the vacuolar lumen.

To summarise, vacuoles are affected by heat shock in several ways; a decrease in number, an increase in size and increased luminal pH.

### Nucleus and mitochondria change in size and develop electron dense content throughout heat shock progression

The nucleus was also affected by the increase in temperature (fig. 4A). Surprisingly, a significant shrinkage (11% smaller than before heat shock) occurred 15 min after temperature shift (fig. 4B). The nucleus then recovered the same approximate size as before. Nuclear morphology was also altered and electron dense content (EDC, arrow in fig. 4A) became more prominent as heat shock progressed. The proportion of nuclei containing EDC increased rapidly with the onset of heat shock, coinciding with the decrease in size, but had decreased again slightly by 90 min (fig. 4C).

**Figure 4.**
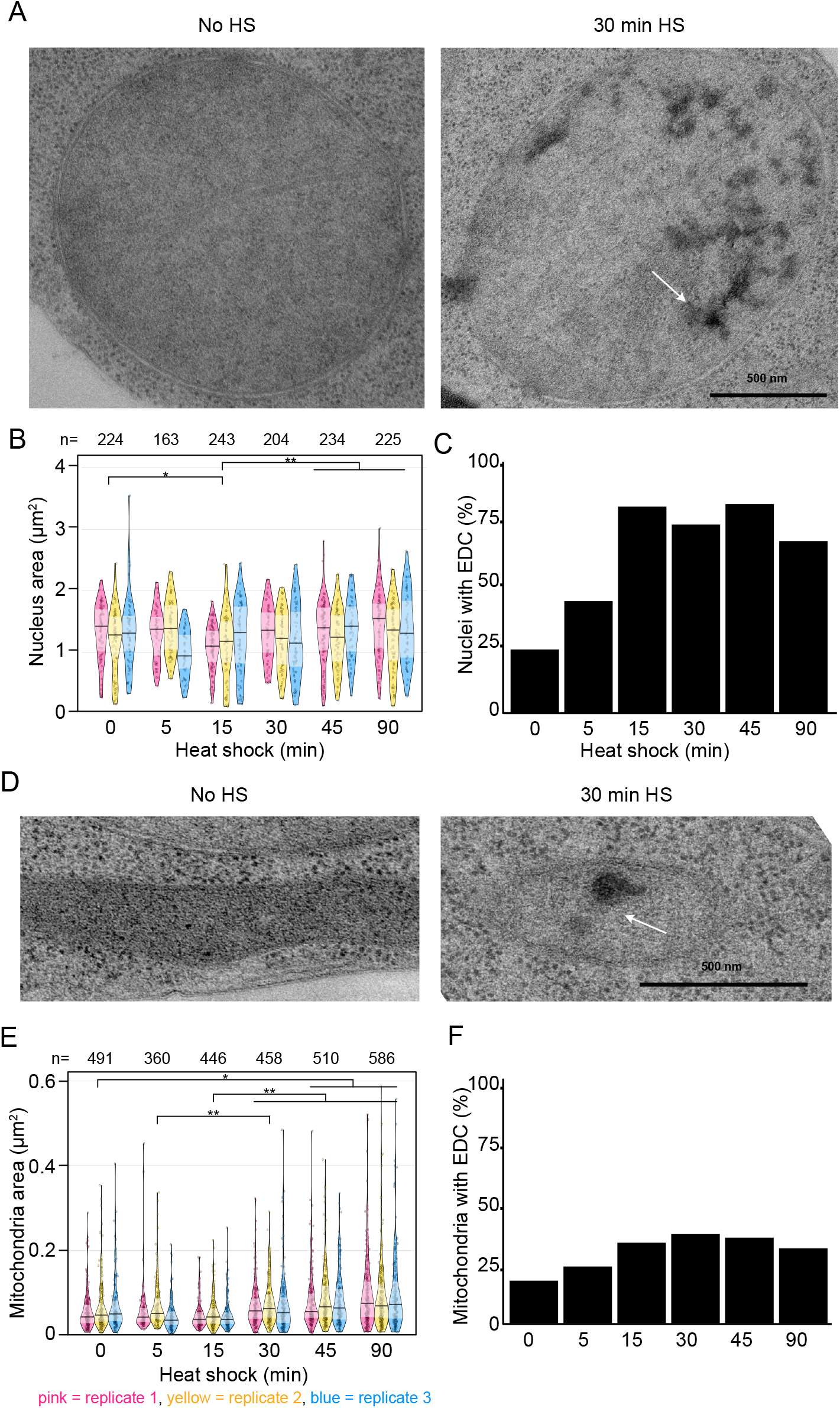
Nucleus and mitochondria change in size and morphology. A) Nucleus morphology before and after 30 min heat shock, as electron dense content (EDC, arrow) appears. B) Nucleus area in electron micrographs of thin sections. The pink, yellow and blue shapes represent the three separate replicates. The width represents the number of measurements within a certain range of values, one point represents one measurement. Solid line is the median and inference bands are the interquartile ranges. C) Proportion of nuclei with EDC throughout heat shock. n=1293 nuclei. D) Mitochondria morphology before and after 30 min heat shock, as EDC (arrow) appears. E) Mitochondria area in electron micrographs of thin sections. On average, the mitochondria increase by 52% over the course of 90 minutes. F) Proportion of mitochondria with EDC throughout heat shock, n=2851 mitochondria. Significances: (*) p < 0.05, (**) p ≤ 0.01.

As one cell section may contain several mitochondria (fig. 4D), 2851 mitochondria were analysed (fig. 2E). Similar to the nucleus, mitochondria were only 75% of their size in untreated cells after 15 minutes of heat shock and significantly different from all time points thereafter. However, eventually a total increase in mitochondrial area of 52% after 90 min was noted. The number of mitochondria stayed relatively constant (fig. S2). The proportion of mitochondria enclosing EDC followed a similar trend to the nucleus, albeit with a less dramatic increase (fig. 4F).

Thus, during heat shock both nuclei and mitochondria vary in size throughout heat shock and also accumulate structures visible as EDC by electron microscopy.

### MVBs and LDs increase in size during heat shock

Two of the smallest cellular organelles, MVBs (fig. 5A) and LDs, displayed the greatest alteration in size over the heat shock time course, with both organelles’ sizes increasing dramatically. The number of MVBs fluctuated during the time course, and after 15 min the cells contained 24% more MVBs than before heat shock (fig. 5B). At the 15 min timepoint MVBs are also significantly different in size from the control (fig. 5C), and after 90 min they had increased in size by 73% (fig. 5C).

**Figure 5.**
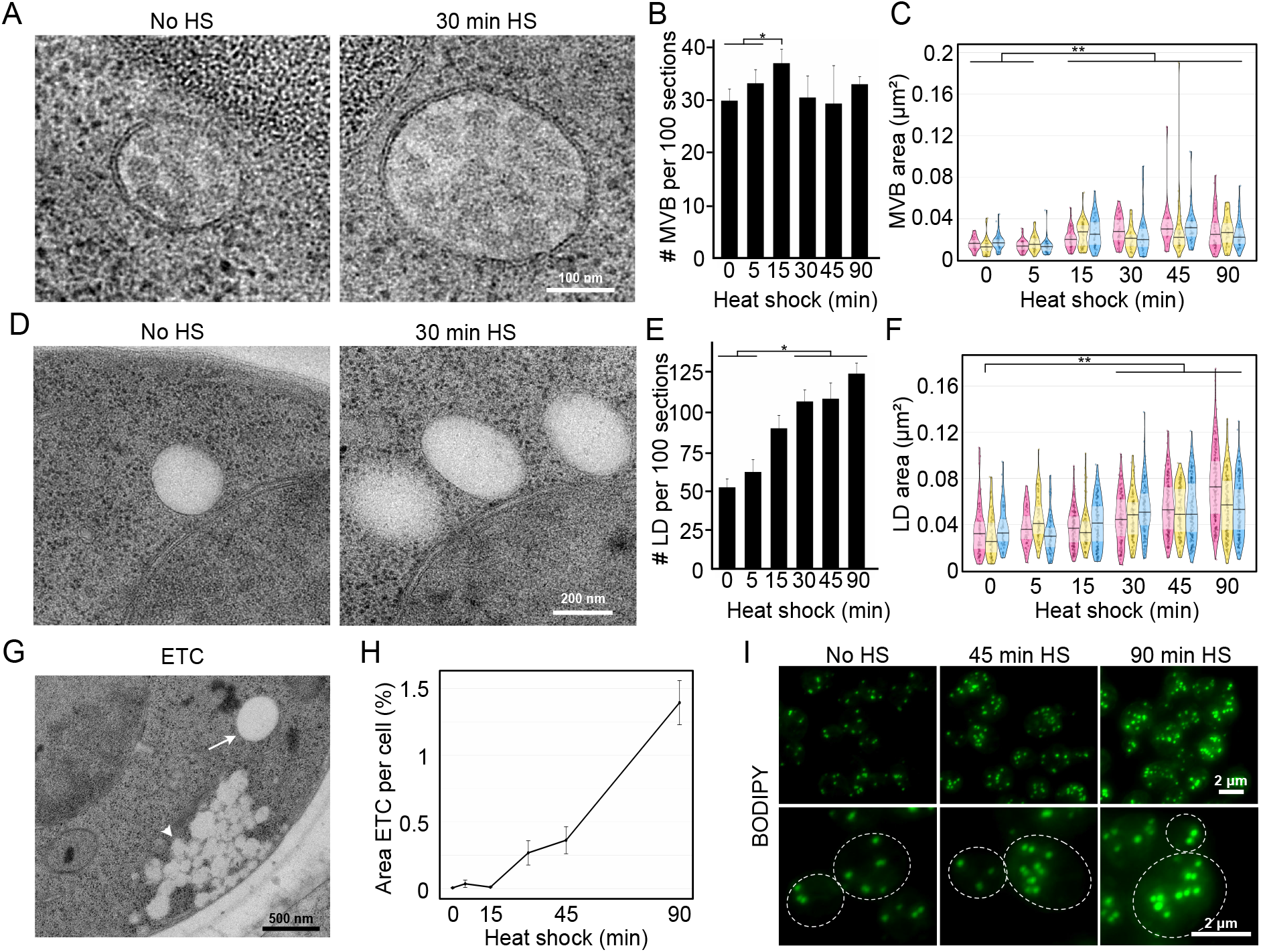
MVBs and LDs increase in size and a new phenotype appears. A) MVB morphology before and after 30 min of heat shock. B) Number of MVBs, averaged over three sets with error bars corresponding to standard error of the mean. C) Area of MVBs, increased by 73% on average. The pink, yellow and blue shapes represent the three separate experiments. The width represents the number of measurements within a certain range of values, one point represents one measurement. Solid line is the median and inference bands are the interquartile ranges. D) LD morphology before and after 30 min of heat shock. E) Number of LDs, increased by factor 2.4, averaged over three sets with error bars corresponding to standard error of the mean. F) Area of LDs, increased by 85% on average. G) Electron translucent clusters (ETC) (arrowhead) next to an LD (arrow). H) Area of the cell section that was covered with electron translucent clusters over time in heat shock. I) Maximum projections of fluorescent images of cells stained with BODIPY for visualization of LDs. Significances: (*) p < 0.05, (**) p ≤ 0.01.

LDs (fig. 5D) also increased dramatically in number, and in cells subjected to 90 min of heat shock they were found 2.4 times as frequently as in the control group (fig. 5E). The first significant difference in size was observed after 30 min, and by 90 min they had increased in size by 85% compared to their equivalent in untreated cells (fig. 5F). In brief, temperature stress leads to a general increase in the size and number of these small organelles.

### Electron translucent clusters appear quickly after heat shock initiation and increase throughout

Previously undescribed grape-like clusters of non-membrane enclosed electron translucent material appeared in heat shock (arrow head; fig. 5G). We named them electron translucent clusters (ETC) and could clearly distinguish them from electron translucent LDs (arrow; fig. 5G), which are larger, clearly delimited from the cytoplasm and never appeared in such large clusters. Also, whilst LDs were most often observed close to the nucleus (fig. 5D and S3), ETC were often found in close proximity to the plasma membrane (fig. 5G and S3). The proportion of the cell area occupied by ETC steadily increased during the heat shock time course (fig. 5H). After 90 min at 38°C, an average of 1.4% of the cell’s cross-sectional area was occupied by ETC, compared to barely having been present before heat shock (when 0.006% of the area was occupied by ETC, n = 361 cells).

To investigate whether the content of ETC and LDs was similar, we stained neutral lipids prominent in LDs (Murphy, 2001) using BODIPY, and compared fluorescence in untreated and heat shocked cells (fig. 5I). The fluorescence increased in a manner that corresponds well with the larger and more common LDs, but was not located near the plasma membrane where the ETC were found in electron micrographs, indicating that the composition of ETC differs to that of LDs. Therefore, throughout the heat shock time course, ETC of an unknown substance increasingly aggregate within the cell near the plasma membrane.

### Physical interaction between vacuole and nucleus increases during heat shock

Not only are organelles of the cell influenced by heat shock as individual components, but also their interactions with each other are affected. Organelles can communicate by forming areas of direct physical contact, called membrane contact sites (MCSs) enabling the exchange of ions, lipids, and signals between the organelles (Gottschling and Nyström, 2017). Although some MCSs are more intensively studied (Pan *et al*., 2000; Kornmann *et al*., 2009; Hönscher *et al*., 2014), MCSs have been identified between almost every cellular organelle (e.g. (Liu *et al*., 2017)). The high-resolution information provided by electron microscopy was used to quantify MCSs between nucleus and vacuole and between vacuole and mitochondria throughout the heat shock time course. An MCS was defined as membranes from two organelles in proximity to each other and not separated by any cytosolic ribosomes and its relative length was calculated in relation to the circumference of the adjoining organelles.

Contact sites between the nucleus and vacuoles (NV, fig. 6A), were not significantly more frequent (fig. 6B) but had increased in length by a total of 57% after 90 min of heat shock (n = 1033 sections, fig. 6C). The number of contact sites between vacuole and mitochondria (VM, fig. 6D) fluctuated strongly, both between time points and sets (fig. 6E). Despite a significant decrease in relative length between 30 and 45 min, when vacuoles are at their largest, the size of VM contact sites did not significantly change (fig. 6F and fig. S4). When multiplying the proportion of sections that contain a contact site with the average absolute length, the surface area for potential interaction between the two organelles can be estimated and shows the same trend as the change in relative length (fig. S4).

**Figure 6.**
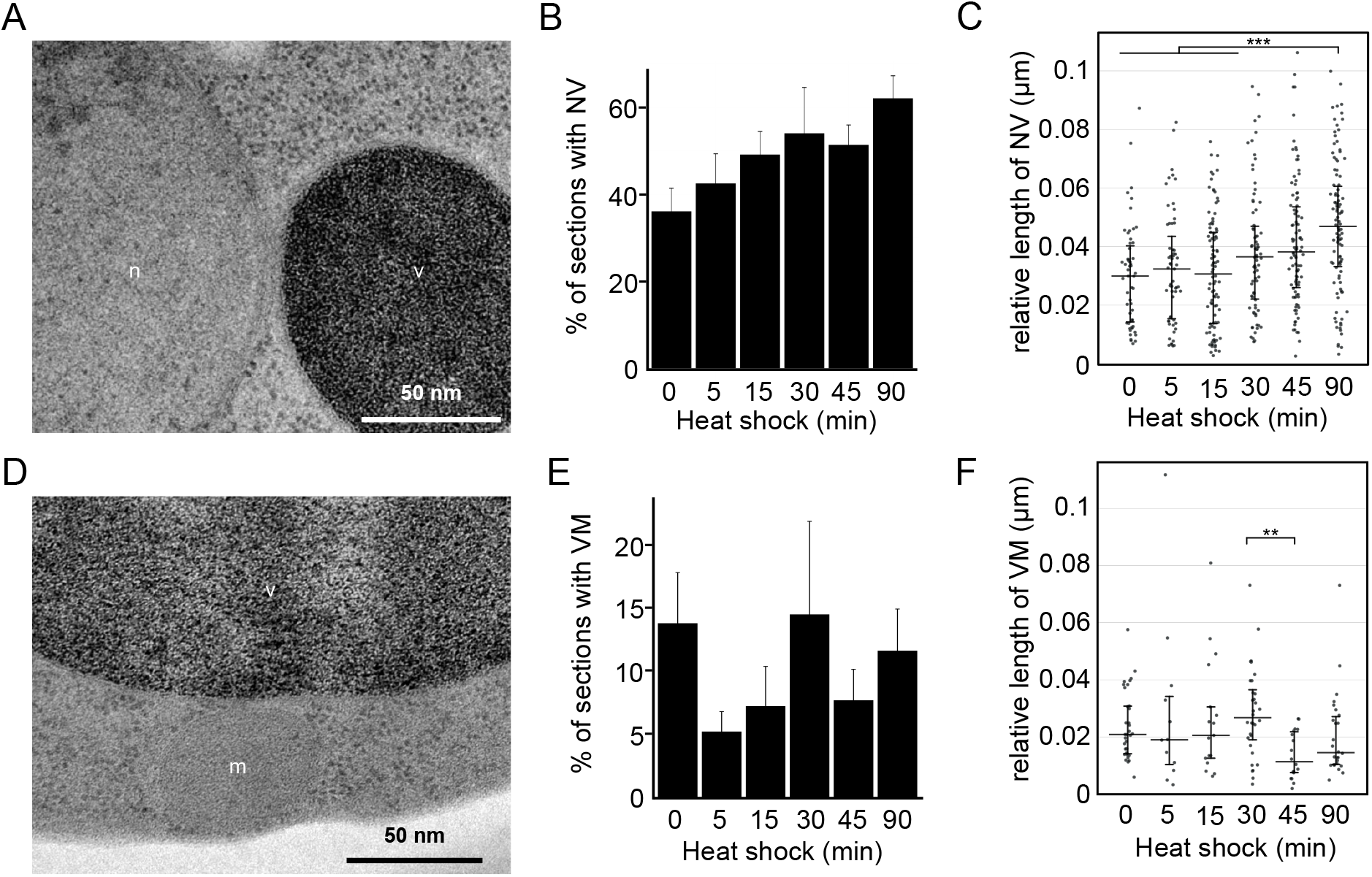
Membrane contact sites are influenced by heat shock. A) Contact site between nucleus and vacuole (NV) B) Percentage of sections containing both nucleus and at least one vacuole with a contact site between the two. C) Length of contact site in relation to circumference of nucleus and vacuole. Black line is the median and error bars the interquartile range. D) Contact site between vacuole and mitochondrion (VM). E) Percentage of sections containing minimum one vacuole and minimum one mitochondrion with a contact site between the two. F) Length of contact site in relation to circumference of vacuole and mitochondria. Black line is the median and error bars the interquartile range. Significances: (**) p ≤ 0.01, (***) ≤ 0.001.

In conclusion, we find that 90-min heat shock appears to specifically increase inter-organellar contact between nuclei and vacuoles but seems to have no to little observable effect on the physical interaction between vacuoles and mitochondria.

## Discussion

Our large-scale electron microscopy approach allows us to both analyse and quantify cellular changes during heat shock in a broad perspective and at high resolution. This showed that mild heat shock affects the size and form of every organelle we studied (fig. 7A), some more severely than others, and some with no previously known roles during heat shock, such as the MVB. It also allowed us to describe a completely new cellular structure – ETC.

**Figure 7.**
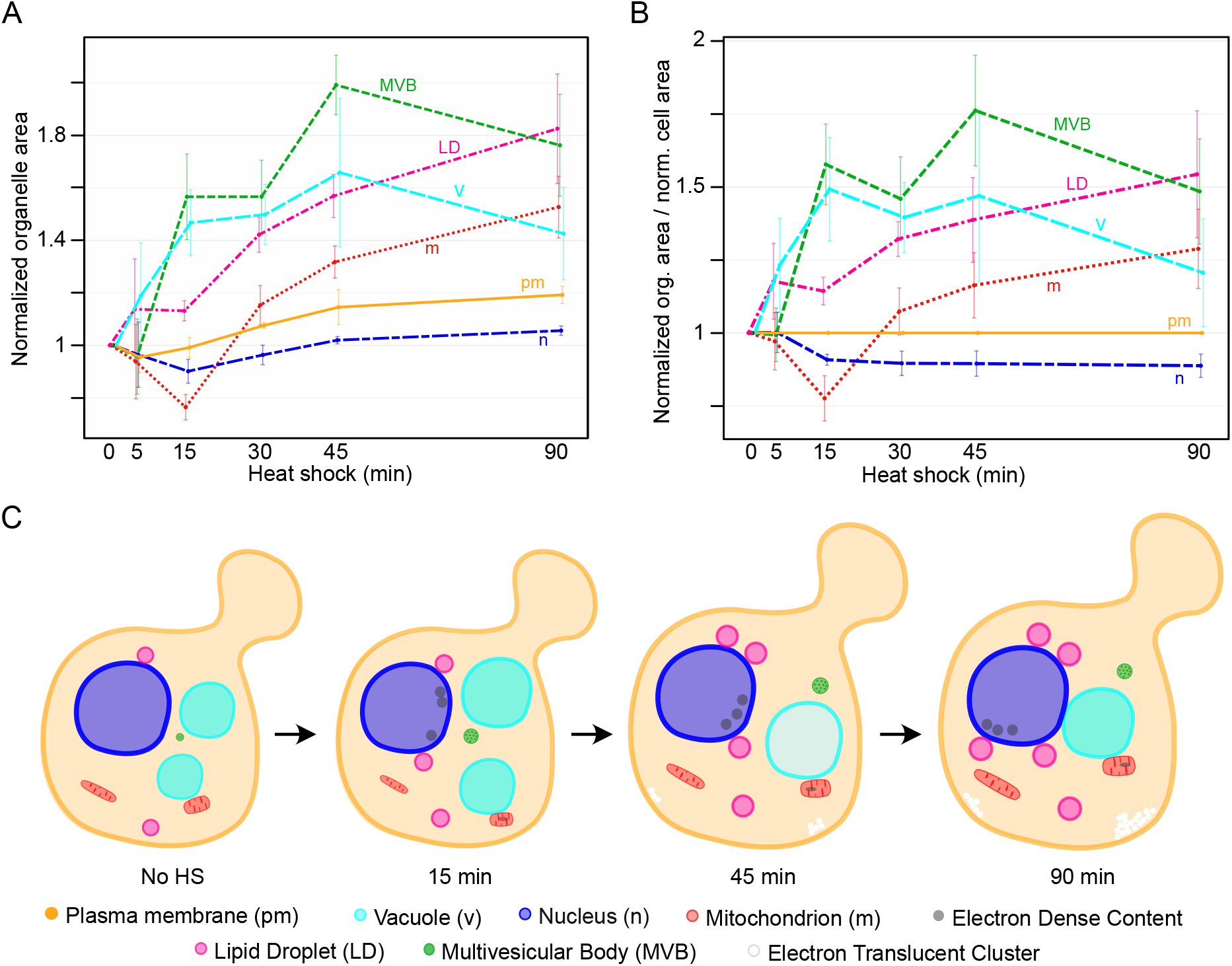
Model of structural changes occurring during mild heat shock. A) Change in organelle area over heat shock with starting size normalized to 1. Points represent the triplicate mean and error bars the standard error of the mean. B) Change in organelle area in relation to cell area at the respective time point with starting sizes normalized to 1. C) Cartoon model of the changes in the whole cell over the course of 90 min.

### A map of large architectural changes to the cell during heat shock reveals dynamic changes in organellar size

One could get the impression that the cell and its organelles respond to heat stress with a general size increase. This could be an effort of the cell to prevent macromolecular crowding, which influences diffusion within the cell as well as protein folding and aggregation (Ellis, 2001). To observe other intracellular interactions, organelle sizes were normalised and plotted in relation to the normalised cell area (fig. 7B), and their respective adaptations became clearer. The organelles’ change in size could be matched up in pairs: cell-nucleus, mitochondria-LD and vacuole-MVB. After the 15 min timepoint both the cell and the nucleus have the same rate of increase in size. Vacuoles and MVBs both rapidly increase in size within the first 15 min and show similar rates of change in size throughout the rest of the time course. Mitochondria and LDs also have similar rates of increase in size starting the 15 min timepoint, revealing dynamic interaction between the two organelles. Indeed, LDs have previously been observed to serve as energy source for mitochondria during nutrient stress (Rambold, Cohen and Lippincott-Schwartz, 2015). In summary, this approach revealed a rapid, complex, and organelle-specific structural response to heat shock (fig. 7C).

### ETC are triggered by heat stress but are unlike LDs in their composition

To our knowledge there are no previous descriptions of ETC in literature although similar structures are clearly visible in previously published electron micrographs of ts mutants at restrictive temperature (Novick, Field and Schekman, 1980; Poon *et al*., 1999; Marini *et al*., 2020). Here, we can only speculate as to their nature and function: the ETC’s electron translucency is similar to that of LDs, indicating that they potentially are lipid deposits. However, the difference in morphology and the lack of BODIPY staining at the cell periphery speaks against the ETC being composed of the same neutral lipids that are found in LDs (Hsieh *et al*., 2012). Further, the majority of LDs are found in proximity to the nucleus, both in untreated cells and after 90 min heat shock (fig. S3). By contrast, other membrane-less structures that are known to accumulate during stress, such as p-bodies and stress granules, would be visible in electron micrographs as electron-dense structures as they contain mRNA which is stained dark by most electron microscopy preparation protocols (Cougot *et al*., 2012).

As opposed to other cellular components, throughout the heat shock time course ETC continued to steeply increase, accumulating near the plasma membrane. A possible explanation for this localisation could be related to the membrane’s fluidity, which is known to increase under stress, especially heat-stress, in various organisms (Dynlacht and Fox, 1992; Lloyd *et al*., 1993; Mejía, Gómez-Eichelmann and Fernández, 1995; Török *et al*., 2014), and has been suggested to initiate the heat shock protein response (Balogh *et al*., 2005). A more fluid membrane, which has been observed to correlate with higher permeability (Wilkes *et al*., 1989), could allow material, such as extracellular fluid or components of the cell wall, to be released into the cytoplasm and observed as ETC in electron micrographs. Furthermore, unsaturated lipids may be deposited nearby to regulate membrane rigidity in higher temperatures.

Alternatively, it has been shown that the sugar trehalose accumulates in cells as a cytoprotectant during stress (Hughes and Gottschling, 2012) and the majority of carbohydrates, just like neutral lipids, are not fixed and stained by our preparation protocol, resulting in electron translucency. It is thus also possible that the ETC could be an accumulation of these carbohydrates.

### MVBs are involved in the cell’s heat shock response

MVBs are involved in the transport of membranous and cytoplasmic content but have no documented roles in the heat shock response to date. Although the number of MVBs remains constant throughout heat shock, their size increases significantly. This could mean that although their synthesis is not affected, the activity of existing MVBs is nevertheless up-regulated. Because intraluminal vesicles formed from the MVB membrane do not only internalize membrane and membrane proteins, but also cytoplasmic content, they offer a proteasome-independent degradation of both membrane- and cytosolic proteins. Thus, their large increase in size throughout heat shock could indicate a need to support the cell’s degradation and sorting machinery as MVBs transport ubiquitinated cargo (Piper and Katzmann, 2010). Interestingly, the increase in size of MVBs and of vacuoles follows a similar path (fig. 2E and 2F), supporting their potential influence on each other’s activity.

### Vacuolar pH is affected by heat stress

Besides an increase in size by almost half their original size, one of the most immediately noticeable changes are the difference in the morphology of the vacuoles. Due to the nature of the preparation protocol, all sets show fluctuations in the electron density of vacuoles, however set 3 is an outlier in its overall high numbers of electron-dense vacuoles. Our results strongly suggest that the different vacuolar electron densities reflect its internal pH and that the uranyl acetate stain can act as a potential pH probe in electron micrographs.

We see that heat stress affects the vacuolar pH in a similar manner that ageing does. In electron micrographs of dividing cells, the daughter had either higher or equal vacuolar electron density when compared to its mother. This is in line with observations that the pH of the vacuole is increased in the mother already after one division (Hughes and Gottschling, 2012). It may seem that deacidification of vacuolar pH during heat stress stems from an export of protons to the cytoplasm to cause the acidification necessary for the induction of HSF1, the main regulator gene of the heat shock response. However, it has been suggested that cells instead depend on the import of extracellular protons for acidification (Triandafillou *et al*., 2019). Observations of vacuolar invaginations, indentation and cytoplasmic vesicles (fig. S1 and (Ishii *et al*., 2018)) may indicate an indirect support of cytoplasmic acidification by increasing the proportion of imported protons relative to cytoplasmic content. The cause and mechanism for vacuolar deacidification remain unclear but the resulting impairment in function (Hughes and Gottschling, 2012) offers a potential explanation for the increase in the size of MVBs due to a block of trafficking to the vacuole.

### MCSs’ response is specific and not directly proportional to organelle size

It appears that the nuclear-vacuolar interaction is of particular importance for the heat shock response since the increase in size of contact sites between the nucleus and vacuoles during heat shock is much higher than would be accounted for by only the enlargement of the corresponding organelles. NV junctions have been shown to be sites for piecemeal microautophagy of the nucleus (Roberts *et al*., 2003) and localisations involved in lipid metabolism (Kohlwein *et al*., 2001; Levine and Munro, 2001).

Similarly, despite an increase in size of both vacuoles and mitochondria, contact sites between the organelles do not increase in size and number, as may be expected. Since vacuoles and mitochondria are closely linked and can form MCS, and vacuolar deacidification contributes to mitochondrial deterioration (Hughes and Gottschling, 2012), it is possible that keeping the number and size of contact sites constant prevents excessive damage to the cell during heat stress. Overall, this shows that there is a specificity in the regulation of the heat shock-induced response of organelle contact sites, the investigation of which has the potential to reveal much about cellular interconnectivity and metabolism under temperature stress. Due to the large size of vacuoles in comparison to the observed microscopy sections, contact sites with vacuoles may be underrepresented in quantification.

### A holistic approach relevant across multiple fields of research

It is valuable to track global cellular changes occurring during heat shock to better understand how the stress response is coordinated. It is also important to be aware of these global changes when for example working with ts mutants where phenotypes attributed to the altered ts allele may instead be a direct or indirect effect of the heat shock itself.

Overall, this study has revealed the ways the cell is influenced by heat shock on a morphological level. Changes in structure, number and size of organelles is related to the molecular activity behind those processes and their careful and balanced interplay. This quantification of organelle changes highlights the importance of a holistic approach to answering research questions and creates a map and reference for those interested in reaction and adaptation to stress by eukaryotic cells.

## Materials and Methods

### Yeast strains

Yeast strains used in this study are derivatives of BY4741, here referred to as wt. Deletion strains have the gene of interest replaced by the KanMX cassette (EUROSCARF). *HSP104-GFP* and *VPH1-GFP* are from the GFP-tagged collection (Huh *et al*., 2003). Strains are listed in Table 1.

### Growth conditions

Cells were cultured at 30°C in rich YPD medium to mid-exponential phase. For heat shock, cell cultures were shifted to 38°C for indicated times before further analysis. The 0 min time point refers to the cultures at 30°C, immediately before being exposed to heat shock.

### Fluorescence Microscopy

For vacuolar pH analysis, 1 OD_600_ of logarithmically growing cells were washed once in YPD + 100 mM HEPES pH 7.6 and stained with 50 μM BCECF-AM (ThermoFisher, Waltham MA, USA) dissolved in the same buffer at 30°C for 30 min. Cells were washed twice in pre-warmed 100 mM HEPES pH 7.6 + 2% glucose, resuspended in pre-warmed YPD and allowed to grow at either 30°C or 38°C for 45 min. Cells were washed in pre-warmed 100 mM HEPES pH 7.6 + 2% glucose then immediately imaged.

For visualisation of lipid droplets, approximately 1 OD_600_ of wild type cells were harvested by centrifugation, resuspended in 1 ml PBS, and incubated with 1 μg/ml BODIPY 493/503 (ThermoFisher, Waltham MA, USA) diluted in DMSO for 15 min. Cells were washed once in PBS before imaged.

Hsp104-GFP and Vph1-GFP expressing cells were grown logarithmically, collected by centrifugation, and imaged directly.

Imaging of live yeast cells was performed with a Zeiss Axio Observer Z1 inverted fluorescent microscope equipped with an AxioCam MRm camera (Zeiss, Oberkochen, Germany). Plan Apo 100x oil objective NA:1.4 and the filter set 38 HEeGFP was used.

### Electron Microscopy

Yeast cultures were grown to an OD_600_ of 0.5, control cells were kept at 30°C and treated cells were heat shocked at 38°C. After removal of medium by filtering through a 0.22 μm filter, the cells were scraped off the filter membrane with a toothpick, transferred to an aluminium carrier and high-pressure frozen in a Wohlwend Compact 3 (M. Wohlwend GmbH, Sennwald, Switzerland). Freeze substitution was performed in a Leica AFS2 (Leica Microsystems GmbH, Wetzlar, Germany) with an incubation in 2% uranyl acetate in acetone (UA; SPI Supplies, West Chester PA, USA) for 1h, rinsing in acetone and embedding in Lowicryl HM20 resin (Polysciences, Warrington PA, USA) (Hawes *et al*., 2007). Polymerised resin blocks were sectioned to 70 nm thickness using a Reichert Ultracut S (Reichert, Vienna, Austria) and placed on Formvar-coated 200-mesh or slot copper grids. Grids were contrast-stained using a 2% aqueous UA solution for 5 min and Reynold’s lead citrate (Reynolds, 1963) 1 min. Samples were imaged at 120kV on a FEI Tecnai G2 Spirit with an FEI Ceta 16M camera (4k x 4k) (Thermo Fisher Scientific, Waltham MA, USA) and a pixel size of 1.1 nm for a magnification of 9300x.

### Image Analysis and Quantification

Number of vacuoles per cell was determined in fluorescence microscopy images of Vph1-GFP cells by manual counting of ≥200 cells per replicate and time point in Fiji (Schindelin *et al*., 2012). Vacuolar area was quantified for ≥200 vacuoles per replicate and time point using the freehand area tool in Fiji on maximum projections of microscopy images of Vph1-GFP cells.

Electron micrographs were quantified using the program IMOD (Kremer, Mastronarde and McIntosh, 1996), by drawing outlines of the structures of interest. Structures such as vacuoles, mitochondria and nuclei were also classified according to morphology. In total, 2143 micrographs were analysed for the time course, corresponding to a full-cell volume of 37 cells, assuming a cell diameter of 4 μm. The length of contact sites was put in relation to the circumference of the respective organelles in contact.

Vacuolar acidity was quantified using Fiji by measuring BCECF-AM mean fluorescence intensity of ≥30 vacuoles per condition from three independent experiments respectively.

### Statistical analysis

After the measurements were obtained from the models, replicates 1, 2, and 3 were treated as one batch. Normality for each timepoint and organelle was checked using a *Shapiro-Wilk-test*. Where possible (cell and MVB area), measurement distribution was normalised using a Box-Cox transform (Manolson *et al*., 1992) (l=0.63 and l=0.4), equal variances confirmed with *Bartlett’s test* and one-way ANOVA performed (cell area). In the case of unequal variances, pairwise t-tests using non-pooled standard deviation were performed (MVB area). Pairwise group comparisons were performed using the R package *emmmeans* (Lenth *et al*., 2020). Non-normally distributed data that could not be normalised was analysed with *Dunn’s test*. All p-values were adjusted according to *Holm* (Holm, 1979).

Where significant differences between replicates 1, 2, and 3 were found, sets were analysed individually and the most conservative p-values shown, these instances are mitochondrial and LD area. Scatter plots show non-transformed data to allow more intuitive interpretation of area values.

## Acknowledgements

The authors would like to thank Mattia Donà, Claes Andreasson, Sabrina Büttner, and Martin Ott for insightful and helpful scientific discussions.

## Data Availability

No publicly available data sets were generated.

## Supplementary figures

**Figure S1.**
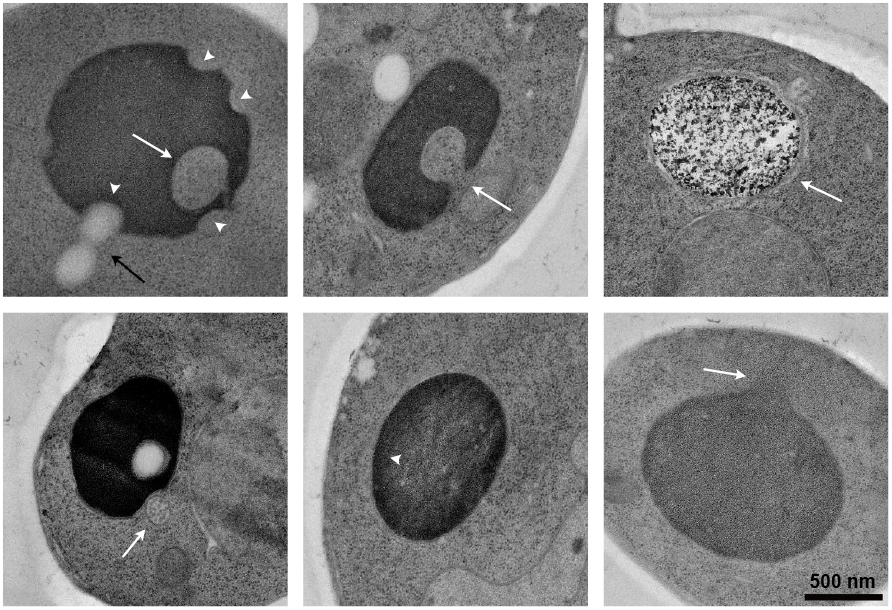
Gallery of vacuolar morphologies in heat shocked cells. (Top) Left: indentations in the vacuolar membrane (arrowheads), LDs (black arrow) and cytoplasmic vesicle contained inside the vacuole (white arrow), Middle: Invagination into vacuole (arrow), Right: Dark precipitates deposited onto the vacuole (arrow). (Bottom) Left: an MVB (arrow) near an indentation of a vacuole, Middle: electron dense ring at the periphery of the vacuole (arrowhead), Right: vacuole seemingly “leaking” content into cytoplasm (arrow).

**Figure S2.**
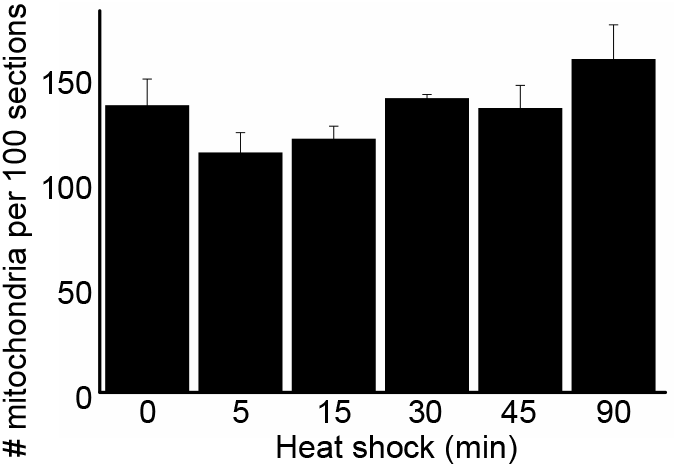
Number of mitochondria does not vary significantly throughout heat shock when observed with EM. A) Number of mitochondria per 100 cell sections in cells subjected to indicated times of HS. 300+ cells were analyzed per time point. Error bars are standard error of the mean.

**Figure S3.**
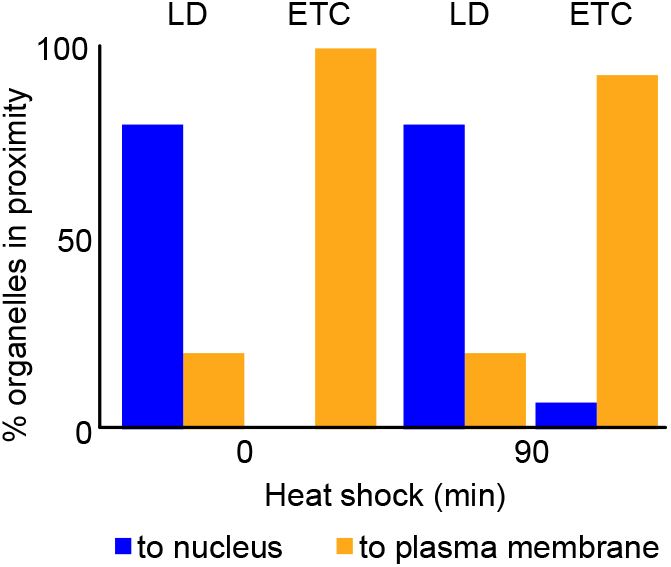
LDs are closer to nucleus and ETC to the plasma membrane. A) Proportion of lipid droplets (LD) and electron-translucent clusters (ETC) respectively closer to the nucleus or plasma membrane before heat shock and after 90 min heat shock. n=50 LD and 22 ETC at 0 min and n=50 LD and 57 ETC at 90 min heat shock.

**Figure S4.**
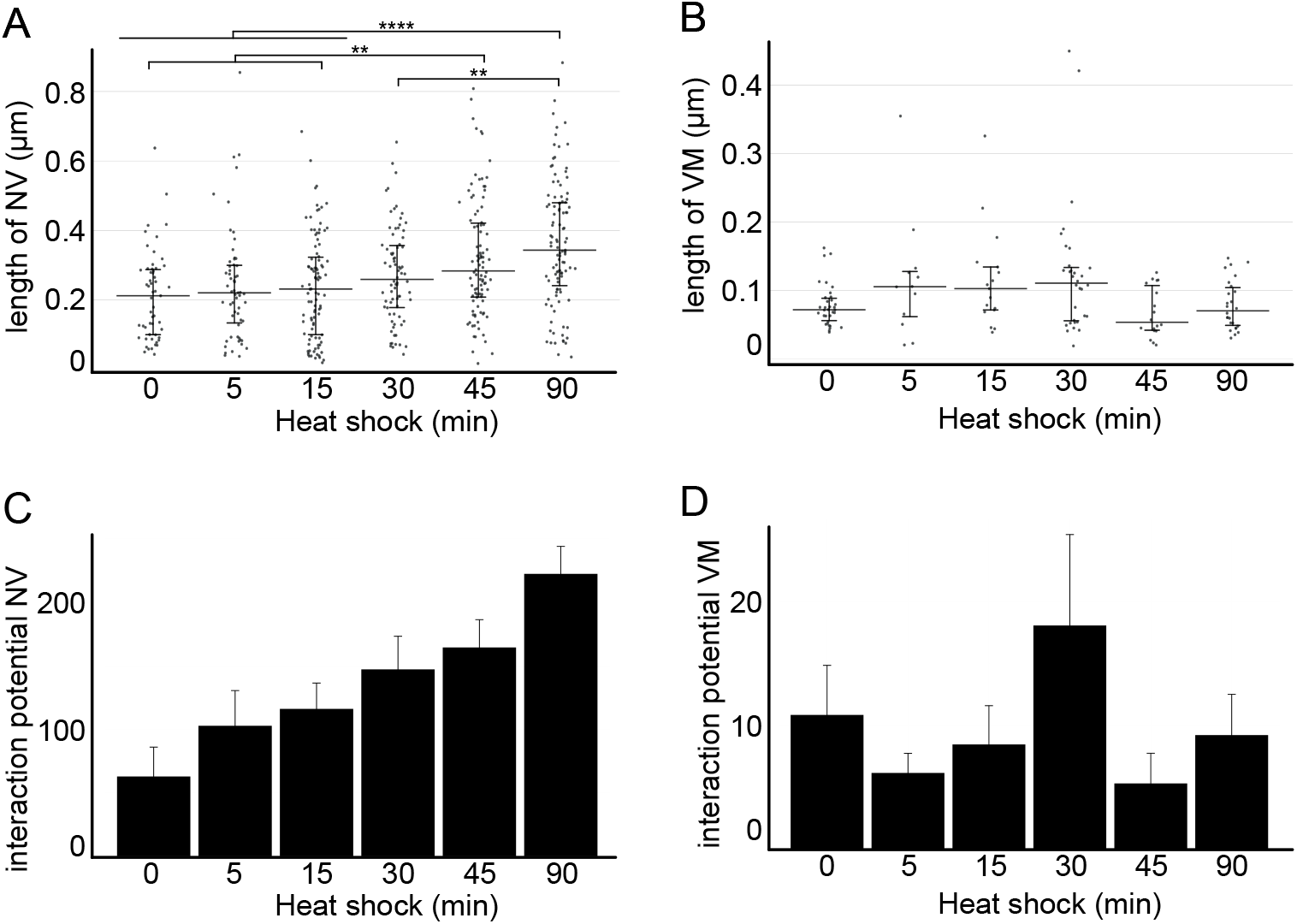
Membrane contact sites are influenced by heat shock. A) Absolute length of contact site between nucleus and vacuole (NV). Black line is the median and error bars the interquartile range. Significances: (**) p ≤ 0.01, (****) ≤ 0.0001. B) Absolute length of contact site between vacuole and mitochondria (VM). Black line is the median and error bars the interquartile range. No significant differences found. C) Interaction potential (arbitrary units) between the nucleus and vacuole, corresponding to the average proportion of sections with contact sites multiplied by the average absolute length in nm. D) Interaction potential (arbitrary units) between the vacuole and mitochondria, corresponding to the average proportion of sections with contact sites multiplied by the average absolute length in nm. Error bars are standard errors of the mean.

